# Molecular memory of periodic thermal stimulation in an immune complex

**DOI:** 10.1101/398966

**Authors:** Razvan C. Stan, Maristela M. de Camargo

## Abstract

Proteins search their vast conformational space in order to attain the native fold and bind productively to relevant biological partners. In particular, most proteins must be able to alternate between at least one active conformational state and back to an inactive conformer, especially for the macromolecules that perform work and need to optimize energy usage. This property may be invoked by a physical stimulus (temperature, radiation) or by a chemical ligand, and may occur through mapping of the protein external environment onto a subset of protein conformers. We have stimulated with temperature cycles two partners of an immune complex before and after assembly, and revealed that properties of the external stimulus (period, phase) are also found in the characteristics of the immune complex (i.e. periodic variations in the binding affinity). These results are important for delineating the bases of molecular memory *ex vivo* and could serve in the optimization of protein based sensors.

---

Organisms respond to environmental cues, such as temperature, by adjusting their physiology and behavior. This property ultimately relies on the ability of the biological molecular machines to construct an internal model of prior interactions with relevant physical and chemical stimuli, in order to predict and adequately respond to a changing environment [1]. This adaptive capacity is manifested by proteins *in vitro*, in cases where phosporilation/dephosphorilation enzymes create cycles of activation/inactivation of the same protein substrate, or when circadian variation in oxygen-carrying ability is promoted in the absence of transcription-translation, through reversible formation of tetramer-dimer complexes [2, 3]. While a cyclical variation in the association and disassociation dynamics between the same or different protein partners generates a corresponding periodic variation in their binding affinity and activity, it has not been established how the system reacts if an external physical stimulus is removed after a few entrainment cycles. This is important for assessing the ability of a biological molecule to anticipate events in its local physical and chemical environment, given the important competitive value this property carries [4]. In particular, we asked whether we can induce in protein partners a periodic change in their binding affinity in response to periodic temperature changes *in vitro*. Underlying the sensitivity of biological systems to temperature is the impact that changes in the thermal energy of the environment has on biochemical and physiological processes, especially in organisms subject to ample temperature fluctuations [5, 6]. Since most of the biochemical activity of a cell consists of short-lived formation of complexes between macromolecules and their ligands, any given binary complex must have a finite useful lifetime. The functional consequences of complex formation are therefore transient and must be tightly regulated in terms of timing of initiation, duration and amplitude of action. We experimentally investigate the theoretical prediction that due to the finite memory of these macromolecules, a selection of the protein conformational states must occur in order to avoid or optimize the energetic cost of erasing irrelevant information caused by stochastic environmental fluctuations [7]. Furthermore, we demonstrate that following application of temperature waveforms, the memory of these periodic external events is manifested in the absence of the original stimulus and may form the basis of a molecular predictive capacity. We have focused here on a monoclonal mouse IgG antibody (MAb k23) relevant for detecting the recombinant antigen, membrane-anchored 19 kDa C-terminal region of the merozoite surface protein 1 (PvMSP1_19_) from the fever-inducing pathogen *Plasmodium vivax* [8]. We measured with isothermal titration calorimetry (ITC) the thermodynamic parameters of their binding after 10-minutes cycles of thermal entrainment at 25◻ and 37◻, prior to complex formation, as shown in Figure 1.

**Figure 1.**
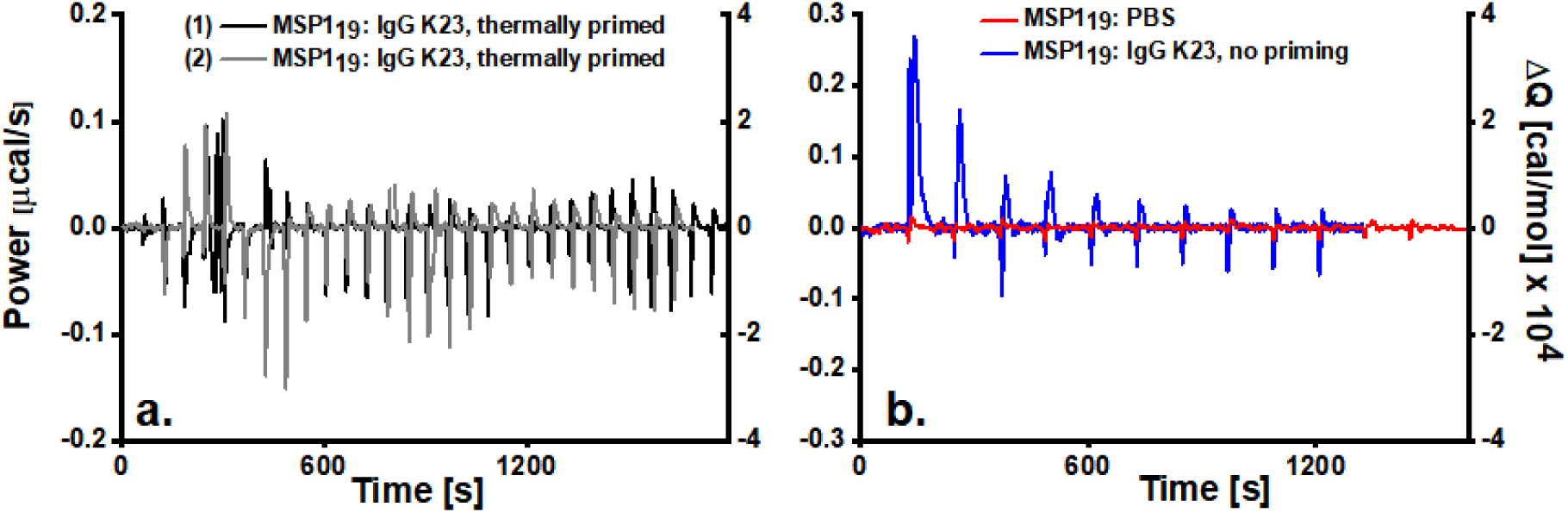
a. Examples of background-subtracted ITC measurements of malarial MSP119 titrated into K_2_3 IgG antibody after prior periodic thermal priming of each protein partner at 25◻ and 37◻. B. Control titrations of MSP1_19_ into PBS buffer (red trace) or with partner IgG K23, without entrainment (blue trace). All measurements performed at 37◻, after 3 cycles at each indicated temperature, last ramp (15 minutes) up from 25◻, before start of titrations, with a pre-equilibration step of 2 minutes.

In contrast to the expected endothermic reaction described by a sigmoid curve [9], the thermograms obtained for the MSP1_19_ titrated into IgG K_2_3 after periodic thermal entrainment revealed a complex sinusoidal pattern composed of both endothermic and exothermic peaks. The presence of intermediate peaks allowed for the determination of the period of oscillations at 530 ± 30 seconds, ≈ 13 % less than the 600-second periods of prior thermal entrainment, a process indicative of underdamped diffusion caused by transient binding events following temperature fluctuations [10]. Importantly, the phase of the oscillation in the binding affinity curve was advanced compared to a fitted harmonic wave [Supporting Information], an observation that may explain a similar phase shift measured *in vitro* with other systems upon temperature increase, on a lengthier circadian basis [11]. We fitted the thermograms to a single-binding site model with 1:1 stoichiometry, for each individual segment of 600 seconds corresponding to the original entrainment periods (Table 1).

**Table 1.** **Thermodynamic parameters obtained by fitting truncated thermograms**

| Segment<br>[minutes] | $K_D$<br>[nM<sup>-1</sup>] | $\Delta H$ [kcal/mol] | $T \cdot \Delta S$<br>[kcal/mol] | $\Delta G$ [kcal/mol] | $k_{on}$<br>[M<sup>-1</sup>s<sup>-1</sup>] · 10<sup>6</sup> | $k_{off}$<br>[s<sup>-1</sup>] | $\tau$<br>[s] |
| --- | --- | --- | --- | --- | --- | --- | --- |
| 0'-10' | 28.4 ± 4.7 | -9.8 ± 2.6 | 9.1 ± 2.3 | -0.6 ± 0.3 | 1.2 ± 0.3 | 0.034 ± 0.005 | 29.4 |
| 10'-20' | 9.8 ± 2.6 | -6.7 ± 2.1 | 4.9 ± 1.2 | -3.2 ± 0.9 | 2.8 ± 0.5 | 0.28 ± 0.3 | 3.5 |
| 20'-30' | 39.4 ± 4.4 | -33.8 ± 4.8 | 37.3 ± 3.7 | -3.5 ± 0.8 | 0.4 ± 0.11 | 0.016 ± 0.003 | 62.5 |
| Control* | 25 ± 3.4 | 17.2 ± 3.1 | 28 ± 2.5 | -10.8 ± 0.6 | 0.29 ± 0.05 | 0.007 ± 0.002 | 142 |

\* MSP1<sub>19</sub> titrated into K<sub>23</sub> IgG, without thermal entrainment treatment.

Prior heat activation in thermal TRP channels is known to profoundly modulate their subsequent temperature responsiveness, causing decreases in the temperature activation threshold while increasing activation time courses [12]. In our system, an important characteristic of the oscillations is the reversal of the enthalpy sign for each truncated thermogram, compared to control titrations, suggestive of net favorable binding. The decrease in entropy in the second segment corresponds to the time frame of entrainment at 25◻ for either complex partner, prior to titrations. We measured for the titrations in the second section a higher binding affinity than either the first (by a factor of ~ 3) or the third segment (by a factor of ~ 4). The increase in entropy in the third segment is significant, suggesting extensive ordering at the binding interface and a conformational selection for each protein partner that presents the most favorable binding interface. Because the affinity alone of a ligand for its binding partner is not a sufficient condition to measure the efficiency and duration of biological action, the lifetime and stability of the binary receptor-ligand complex was also calculated (parameter τ) [13]. This parameter defines the complex stability and thus the effectiveness of interaction between the two protein partners. As with other soluble antibodies, immune complex stability must be optimized, for instance in cases where antigen binding to B-cell receptors occurs through a two-step, induced-fit process prior to endocytosis, requiring sufficiently large residence times for efficient antigen processing [14]. Kinetically, we measure the highest binding affinity in the third segment of the thermograms, accompanied by the lowest k_off_ and thus the largest complex residence time (τ ≈ 1 minute). Interestingly, lowest and highest k_off_ were measured in the second segment, mirroring the thermodynamic observations outlined above.

Following immune complex formation, important information relating to the binding mechanism and the stability of the complex can be obtained using Differential scanning calorimetry (DSC) in isothermal mode, since, unlike ITC, the thermal entrainment cycles and corresponding variations in heat capacity (Cp) are measured *in situ* (Figure 2).

**Figure 2.**
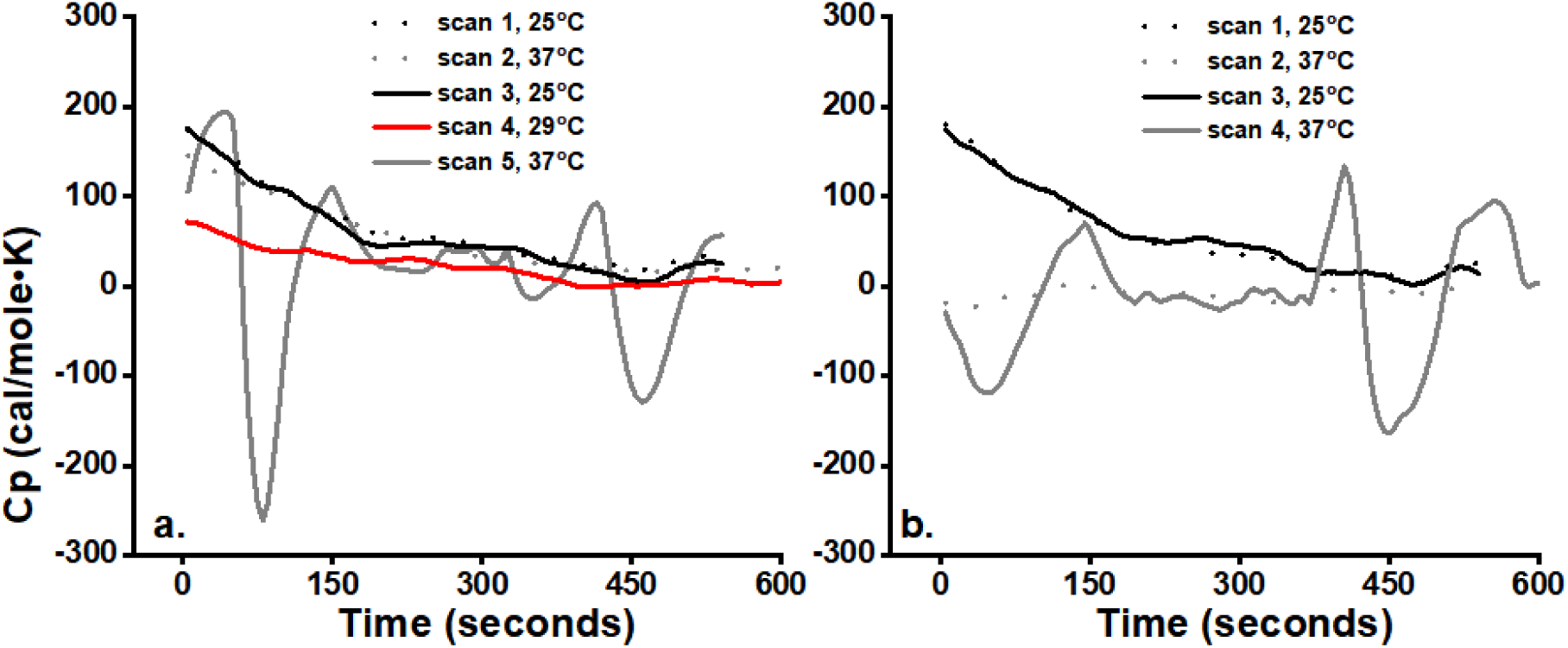
Background-subtracted, normalized DSC scans of equimolar MSP119: IgG K_2_3 (0.5 mg/ml total protein complex). Reconstitution of the immune complex was achieved *in situ* at 37◻ for 30 minutes, prior to temperature ramping. Similar to ITC measurements, following 3 entrainment steps at 25◻ and 37◻ for 10 minutes each, data was collected from scan 1, as indicated in fig. 2.a and 2.b. Pre-equilibration time at 5 minutes, before data scan collection.

Most DSC curves commence with an endothermic effect, indicating that a process had started before it could be monitored, a time limitation imposed by the necessary equilibration period for heating/cooling the samples [Supporting Information]. We measured disruptions of the immune complex accompanied by an increase in heat capacity (positive Cp), followed by a subsequent decrease, indicating immune complex reformation, as observed with other systems [15]. Endothermic and exothermic peaks from the DSC scans were modeled separately with a bigaussian function, whose fitting parameters are shown in Table 2:

**Table 2.** **Fitting parameters for the averaged DSC scans**

| Peak | Peak height [kcal/mole·K] | Peak center [seconds] | Peak width [seconds] |
| --- | --- | --- | --- |
| 1 | 176 ± 44 | 53 ± 12 | 27 ± 12 |
| 2 | -309 ± 21 | 110 ± 31 | 33 ± 14 |
| 3 | 79 ± 17 | 139 ± 22 | 31 ± 13 |
| 4 | 82 ± 23 | 435 ± 29 | 52 ± 18 |
| 5 | -173 ± 31 | 470 ± 34 | 34 ± 12 |

Similar to ITC measurements, we measured autonomous fluctuations in heat capacity changes of the immune complex, characterized by both a marked decrease in peak amplitudes with time, and by phase shifts, as measured from the position of exothermic peak centers. We calculated a period of 390 ± 20 seconds for averaged DSC scans, based on the peak centres of the exothermic oscillations, a value close to the oscillation period measured in ITC. Furthermore, the peak width of the second re-association step of the protein partners is larger than the corresponding first peak width, an increase in the residence time that we also observed in ITC (≈ 1 minute). Importantly, not all DSC curves contained both exothermic peaks, possibly indicating re-association steps that occur during the experimentally inaccessible pre-equilibration periods (Figure S1.a). We have not detected any periodic features in DSC scans after sample removal, storage at 4 and later measurements without thermal treatment, suggesting the complete loss of initial temperature priming information (Figure S1.b).

Similar to other C-terminal regions of *Plasmodia* MSP1, PvMSP1_19_ consists of two successive EGF-like domains, with each EGF-like domain containing a segment of random coil followed by four antiparallel β-strands [16]. The presence of these disordered segments affords the augmented protein sensitivity to thermal changes [17], manifested by increased absorption of the thermal energy from the solvent that is transferred to the highly thermally-conductive β-sheets [18]. As the temperature is cycled between the two values prior to measurements, either protein exhibits sensitization to temperature, an increase in the protein response to external stimulus, as measured elsewhere for temperature-activated TRP channels [19]. We propose that the periodicity we measured is based on the large enthalpy changes between the applied temperature waveforms, manifest across the titrations. As this property is conducive to hysteretic behavior [17], with higher activation energies resulting in anisotropic protein backbone motions [20], we further propose that their reiterated succession results in the periodic features we measured.

The presence of phenomenological molecular memory here described was previously observed in hysteretic enzymes, whereby the protein slowly responds to changes in the concentration of metabolites [21], binding partners [22], in pH [23] or applied voltage [24] thus providing a time-dependent buffering capacity against stochastic oscillations of signalling partners or physical stimuli. Hysteretic behaviour can also be present when the ligand uncouples from its protein partner at a sufficiently slow rate. We hypothesize that protein conformational isomers populate certain regions in the conformational phase space with a higher probability after the action of various stimuli (temperature, ligands etc), or conversely, that the space between the conformers is not isotropic, such that once the macromolecule has settled in a particular energetic well, it will require a higher activation energy to escape it. Free energy barriers must separate the distinct internal memory states of a system, absent which it will randomly fluctuate from one state to another under the influence of thermal fluctuations. However, the periodic optimization in the thermodynamic and kinetic binding parameters we observed, coupled to the increase in k_off_ rates, suggests that a subpopulation of protein conformers is still present for a specific stimulus, even after the system has been subjected to a new and distinct signaling event. Importantly, entropy increase across the truncated thermograms may indicate a causal entropic forcing of the system that maximizes the span of future accessible pathways to stimuli, and thus its adaptive capacity, as theoretically predicted [25]. Series of coupled protein oscillators are also believed to be responsible for the mnemonic and intelligent behavior of single cell eukaryotes following cyclic stimulation [26]. The potential to retain or revisit *in vitro* specific protein conformational states, regularly and at different timescales, after periodic external stimulation, remains thus an area to be explored. Whether this property can be extended to other scenarios, by example through reversibly creating binary and ternary complexes of competing ligands and repeatedly measuring their binding affinity before and after complex formations, will be important for dissecting the *in vitro* the mechanisms of molecular memory.

## Supporting Information Available

Experimental procedures, DSC scans and ITC thermograms are included in the supporting information.

